# Cross-species comparison of pregnancy-induced GDF15

**DOI:** 10.1101/2023.06.19.545552

**Authors:** Anders Bue Klein, Pablo Ranea-Robles, Trine Sand Nicolaisen, Claudia Gil, Kornelia Johann, Júlia Prats Quesada, Nina Pistolevij, Kathrine V.R. Hviid, Line F. Olsen, Simone M. Offersen, Jørn Wulff Helge, Henriette Svarre-Nielsen, Jaco Bakker, Maximillian Kleinert, Christoffer Clemmensen

## Abstract

GDF15 (growth differentiation factor 15) is a stress-induced cytokine. Although the exact physiological function of GDF15 is not yet fully comprehended, the significant elevation of circulating GDF15 levels during gestation suggests a potential role for this hormone in pregnancy. This is corroborated by genetic association studies in which GDF15 and the GDF15 receptor, GDNF Family Receptor Alpha Like (GFRAL) have been linked to morning sickness and hyperemesis gravidarum (HG) in humans. Here, we studied GDF15 biology during pregnancy in mice, rats, macaques, and humans. In contrast to macaques and humans, mice and rats exhibited an underwhelming induction in plasma GDF15 levels in response to pregnancy (∼75-fold increase in macaques vs. ∼2-fold increase in rodents). The changes in circulating GDF15 levels were corroborated by the magnitude of *Gdf15* mRNA and GDF15 protein expression in placentae from mice, rats, and macaques. These species-specific findings may help guide future studies focusing on GDF15 in pregnancy and on the evaluation of pharmacological strategies to interfere with GDF15-GFRAL signaling to treat severe nausea and HG.

## Introduction

GDF15 is a stress-induced cytokine that was initially recognized as an autocrine regulator of macrophage activation^1^. Following the discovery that pharmacological GDF15 has weight-lowering properties in preclinical models^2-5^, the search for an endogenous role for GDF15 in energy metabolism accelerated^6^. This includes work from several labs, including our own, demonstrating that GDF15 is an exercise-regulated hormone^7-13^. Although prolonged exercise can increase endogenous GDF15 substantially (∼5-fold induction in plasma), we have yet to understand the physiological consequences of this increase. By contrast, it is well established that recombinant GDF15 (rGDF15) can lower food intake but also that it induces aversion^10,14^. To date, several studies have reported dose-dependent aversion to rGDF15 in rodents (summarized in^15^), and have also demonstrated that rGDF15 promotes emesis and that the vomiting correlates with the reduced energy intake in the musk shrew animal model (*Suncus marinus*, a non-rodent mammal)^16^. Accordingly, it has been proposed that GDF15 is a conserved defense signal serving to elicit toxin avoidance^17^.

Of note, pregnancy is the only known physiological condition where endogenous GDF15 levels are comparable to the pharmacological doses required to cause aversion and emesis^14,18,19^. During gestation, GDF15 levels can rise up to 200-fold peaking at the end of 3^rd^ trimester^20^. GDF15 is primarily produced and secreted from the extravillous trophoblasts of the placenta but is also highly expressed in decidual stromal cells^21,22^. Given the aversive effects of rGDF15 and because nausea and vomiting affect ∼70% of pregnant women, it has been suggested that this increase in circulating GDF15 governs nausea and vomiting during pregnancy. This is supported by genetic studies linking common variants in GDF15 and GFRAL genes to hyperemesis gravidarum (HG)^23-25^, a condition that has been estimated to affect up to 2% of all pregnancies^26^.

Animal models with loss-of-function mutations have proven to be invaluable tools for investigating causal pathways in physiology^27^. This is also the case for the GDF15-GFRAL pathway for which global and tissue-specific GDF15 knockout (KO) mice and GFRAL KO mice have been used to parse the role of GDF15 in mediating the effects of a number of drugs, including anti-diabetic drugs^28-31^ and chemotherapeutic agents^4,32^. However, GDF15 regulation during gestation in rodents have not been assessed and accordingly it remains unknown whether rodent models are applicable to study the physiological role of GDF15 during pregnancy. In the present study, we address this by investigating placental and circulating GDF15 in mice, rats, macaques, and humans.

## Methods

### Animals

#### Rodents

C57BL/6J female mice and Sprague-Dawley female rats were obtained from Janvier-Labs (Le Genest-Saint-Isle, France). The female GDF15 KO and WT littermate mice were generated as described previously^29^. Animals were group housed and all experiments were done at 22°C with a 12:12-hr light-dark cycle. Animals had free access to water and chow diet throughout the experiments. Animals were time-mated and on day 14 of the gestation period, the dam was euthanized, and embryos removed and placentae snap-frozen using dry ice. Similarly, kidneys and livers were dissected out and snap-frozen using dry ice. Blood samples from the dams were collected in EDTA-coated tubes and centrifuged (5000 x g; 10 min) and plasma collected and stored at -20°C until further analysis. All rodent experiments were approved by the Danish Animal Experimentation Inspectorate (2018-15-0201-01457) or by the LAVG Brandenburg, Germany (2347-14-2021).

#### Humans

For measurement of GDF15 in non-pregnant women, blood samples from Frandsen et al were obtained^33^. Initially, subjects received written and oral information about the study and signed an informed consent form. Blood samples were obtained from an antecubital vein and collected in precooled vacutainers (Vacutainer BD). The blood samples were immediately centrifuged at 4000 rpm for 10 minutes at 4°C and the plasma fraction stored at –80C for later analysis. The original study^33^ was approved by the Ethics Committee in Copenhagen (H-17024872) and conducted in accordance with the last revised ethics guidelines of the Declaration of Helsinki.

For measurement of GDF15 in pregnant women blood samples were obtained from the PREGCO cohort^34,35^. Included samples are from women attending their second trimester ultrasound at the Obstetrics & Gynecology department at Copenhagen University Hospital Hvidovre. Participants received written and oral information about the study and signed an informed consent form. Blood samples were obtained from an antecubital vein and collected in vacutainers (Vacutainer BD). The blood samples were centrifuged at 3350 rpm for 10 minutes at 20°C (Centrifuge Eppendorf 5804R). Serum samples were analyzed for SARS-CoV-2 antibodies, using YHLO’s iFlash 1800 and SARS-Cov-2 IgM/IgG kits. IgG antibody level of 10.00 AU/mL or higher was considered positive and less than 10.00 AU/mL as negative. After analysis serum was stored at –80C. The PREGCO study was approved by the scientific ethics committee of the capital region of Denmark (ref. no. H-20022647).

#### Macaques

All rhesus monkeys (*Macaca mulatta*) were housed at the Biomedical Primate Research Centre (BPRC, Rijswijk, The Netherlands). The BPRC houses an outbred breeding colony, consisting of approximately 800 rhesus macaques. All animals lived with their offspring in naturalistic family groups of approximately 15-30 individuals with an age range from newborn to 30 years old. Animals were housed in enriched enclosures with permanent accessible outdoor compartments. For indoor enclosures, the room temperature was maintained at 20 ± 2°C, a room ventilation rate of six air changes per hour, and with a 12:12-h light:dark cycle (lights on, 7.00 a.m.–7.00 p.m.). The animals were fed commercial monkey pellets (Ssniff, Soest, Germany) supplemented with limited amounts of fruit, vegetables, or grain mixtures. Enrichment containing food was provided every other week. Municipal water was offered *ad libitum* by water nipples. Physical examinations were routinely performed yearly, including a thorough physical examination, a tuberculosis screening test, complete blood count (CBC) and serum biochemistry. All animals were housed in accordance with Dutch law and international ethical and scientific standards and guidelines (EU Directive 63/2010). Any procedures and husbandry were compliant with the above standards and legislation. The animal care at BPRC is in accordance with programs accredited by *AAALAC International*. Blood samples were collected during the annual health evaluations in April and May 2021. No animal was sampled solely for the purpose of this study.

All blood samplings were conducted during between 8.30-12 a.m. Each monkey had last been fed on the previous midday, water was *ad libitum* available. Animals were sedated with ketamine (10 mg/kg ketamine hydrochloride (Alfasan Nederland BV, Woerden, Netherlands) combined with medetomidine hydrochloride (0.05 mg/kg (Sedastart; AST Farma B.V., Oudewater, Netherlands), both administrated intramuscularly. Approximately 2.5 mL blood was collected from a femoral vein by using a 20-gauge Vacuette needle and hub (Greiner Bio-One GmbH, Kremsmünster, Austria) by qualified caretakers. After completing all health surveillance procedures, upon return to their home cage, atipamezole hydrochloride (Sedastop, ASTFarma B.V., Oudewater, Netherlands, 5 mg/ml, 0.25 mg/kg) was administrated intramuscular to antagonize medetomidine. Samples were collected in plain tubes and allowed to clot. Samples were centrifuged at 3000 rpm for 10 min. Serum was prepared from the blood samples and after routine clinical chemistry for health surveillance the leftovers were stored at –80°C until analyzing for GDF15.

All animals underwent a transabdominal uterine ultrasound evaluation using an ACUSON Juniper ultrasound system with a micro-convex 11M3 3.5-11.0 MHz transducer (Siemens Healthcare GmbH, Erlangen, Germany) for pregnancy determination. All ultrasound evaluations were performed and analyzed by experienced veterinarians. The exact parturition dates were obtained in retrospect from the electronic health and medical records of those animals. All animals scored pregnant on physical examination with ultrasound delivered and no not pregnant monkeys delivered, nor abortions were observed. In total 15 pregnant and 14 age-matched female non-pregnant (control) animals were included in this study. All included females were mature and free of signs of clinical disease on the moment of sampling.In case animal caretakers observed a placenta in an animal enclosure, the placenta was collected and stored in -20°C untill further analyses. Kidney and liver were collected from four males and one female macaque. The tissues were stored at -80°C after obtaining. The organs were obtained from animals in the colony that were euthanized (pentobarbital, 70 mg/kg intracardially) for ethical reasons. No animals were euthanized for the purpose of this study.

#### ELISA

GDF15 was measured in human plasman and serum, and macaque serum using the Quantikine ELISA Human GDF-15 Immunoassay (ELISA, R&D systems, catalog no. DGD150). In mice and rats, plasma samples were analyzed using Quantikine ELISA Mouse GDF15 Immunoassay (ELISA, R&D systems, catalog no. MGD150). The ELISA assays were used according to the protocol provided by the manufacturer.

#### RNA extraction & cDNA synthesis

Placenta, kidney and liver were homogenized in a Trizol reagent (QIAzol Lysis Reagent, Qiagen) using a stainless steel bead (Qiagen) and a TissueLyser LT (Qiagen) for 3 min at 20 Hz. Then, 200 μl chloroform (Sigma-Aldrich) was added, and tubes shaken vigorously for 15 sec and left at RT for 2 min, followed by centrifugation at 4°C for 15 min at 12,000 x g. The aqueous phase was mixed 1:1 with 70% ethanol and further processed using RNeasy Lipid following the instructions provided by the manufacturer. For muscle tissue, the lysis procedure described in the enclosed protocol in the Fibrous Tissue Mini Kit (Qiagen) was followed. After RNA extraction, RNA content was measured using a NanoDrop 2000 (Thermo Fisher) and 500 ng of RNA was converted into cDNA by mixing FS buffer and DTT (Thermo Fisher) with Random Primers (Sigma-Aldrich) and incubated for 3 min at 70°C followed by addition of dNTPs, RNase out, Superscript III (Thermo Fisher) and placed in a thermal cycler for 5 min at 25°C, 60 min at 50°C, 15 min at 70°C, and kept at -20°C until further processing.

#### qPCR

SYBR green qPCR was performed using Precision plus qPCR Mastermix containing SYBR green (Primer Design, #PrecisionPLUS). qPCR was performed in 384-well plates on a Light Cycler 480 Real-Time PCR machine using 2 min preincubation at 95°C followed by 45 cycles of 60 sec at 60°C and melting curves were performed by stepwise increasing the temperature from 60°C to 95°C. Quantification of mRNA expression was performed according to the delta-delta Ct method.

### Statistical analyses

Statistical analyses were performed in GraphPad Prism Version 9. For comparing multiple groups, one- or two-way ANOVA with Bonferroni post hoc multiple comparisons test was used. When comparing two groups, a Student’s t-test was used. Unless otherwise stated, all data are presented as mean ± SEM. P < 0.05 was considered statistically significant.

## Results

To assess if pregnancy affects circulating GDF15 levels in mice, we collected blood from pregnant C57BL/6J mice as well as from non-pregnant control mice throughout gestation (**Figure 1A**). Circulating GDF15 levels rose steadily until day 10, where they plateaued at a ∼2-fold increase over non-pregnant mice until the end of the pregnancy (**Figure 1A**). Similarly, we found a 70% increase in plasma GDF15 levels at day 14 of pregnancy in Sprague-Dawley rats (**Figure 1B**). These findings highlight that in contrast to humans^19,20^, circulating GDF15 levels only modestly increases during pregnancy in rodents. We subsequently investigated the effects of pregnancy on plasma GDF15 in macaques. At midterm (middle of 2^nd^ trimester), serum GDF15 levels were approximately 75-fold increased (**Figure 1C**). To validate the similarity between macaques and humans, we measured circulating GDF15 levels in healthy, non-pregnant women and in healthy pregnant woman in the midterm (middle of 2^nd^ trimester). In agreement with previous studies^20,22^, circulating GDF15 levels were increased to ∼25,000 pg/mL halfway through the gestation period in humans (**Figure 1D**). When comparing circulating GDF15 levels across pregnant species, GDF15 levels increased ∼2-fold in rodents (**Figure 1E**), while circulating GDF15 levels increased ∼75-100-fold in humans and macaques (**Figure 1E**).

**Figure 1.**
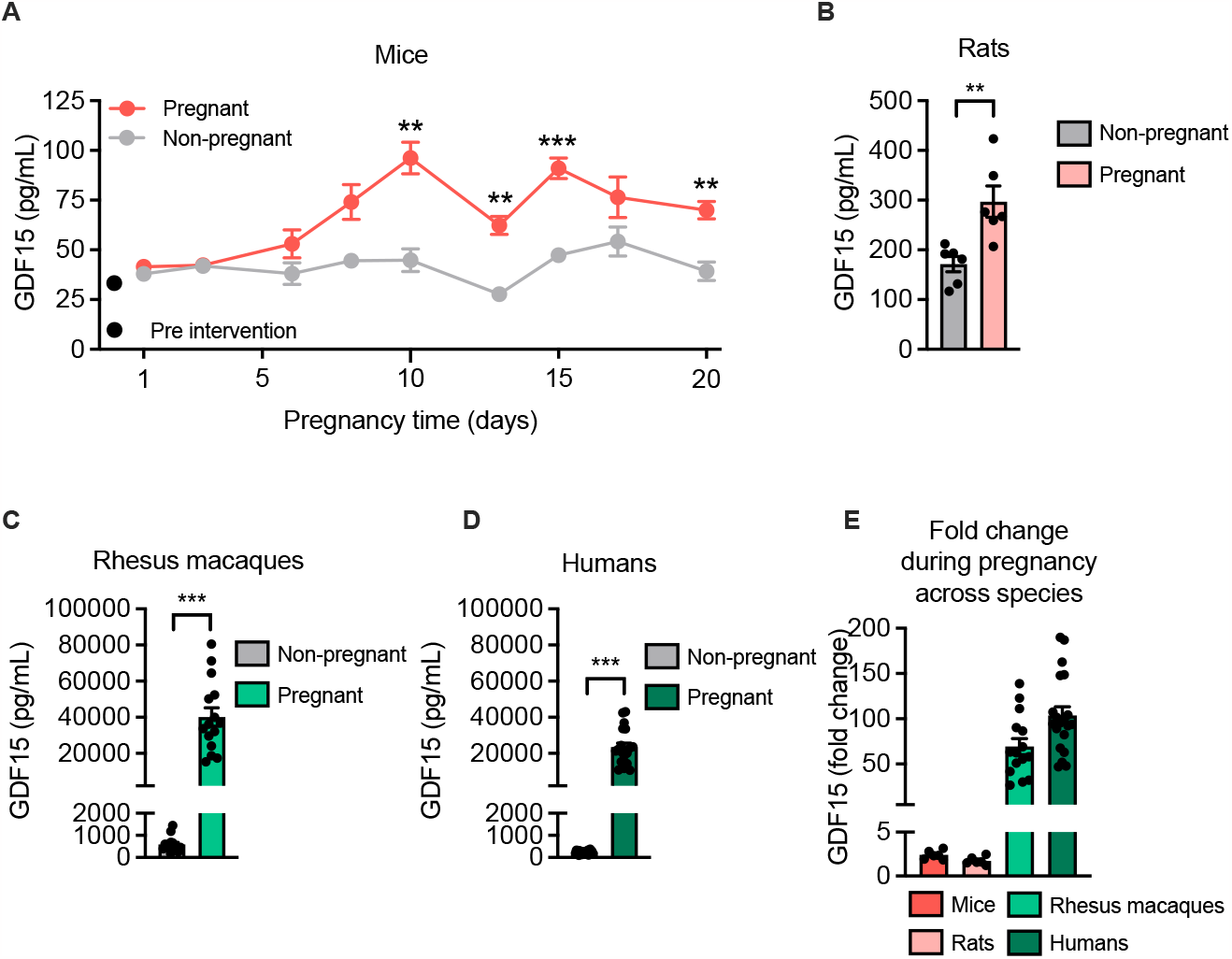
Cross-species comparison of pregnancy-induced GDF15. **A**. Mouse GDF15 plasma levels (pg/mL) in aged matched non-pregnant (n=5) versus pregnant (n=7) C57BL/6J mice at nine different time points during the gestation period. **B**. Rat GDF15 plasma levels (pg/mL) in aged-matched non-pregnant versus pregnant Sprague-Dawley rats (n=6). **C**. Non-human primate GDF15 plasma levels (pg/mL) in aged-matched non-pregnant (n=14) versus pregnant (n=15) macaques. **D**. Human GDF15 plasma levels (pg/mL) in aged-matched non-pregnant (n=22) versus pregnant (n=20) healthy women. **E**. Fold changes in circulating GDF15 levels during pregnancy across species. Data are presented as mean ± SEM. ** p < 0.01, *** p < 0.001.

To further decipher the species differences in GDF15 induction during pregnancy, we measured GDF15 mRNA and protein levels in mice, rats, and macaques. In WT mice, the placental *Gdf15* mRNA levels were similar to liver and kidney mRNA levels (**Figure 2A**). In rats, the same pattern could be observed, with similar *Gdf15* mRNA levels among liver, kidney, and placenta (**Figure 2B**). In contrast, *GDF15* mRNA levels in placenta from macaques were significantly higher than in liver and kidney (**Figure 2C**). When comparing normalized GDF15 mRNA levels in placentas across species, we found that GDF15 mRNA levels were at least 1000-fold higher in monkey placenta when compared to placenta tissue from rodents (**Figure 2D**). The mRNA data were corroborated by protein GDF15 levels assessed by ELISA, demonstrating markedly higher GDF15 protein in placenta than in liver and kidney in macaques (**Figure 2E**). To investigate whether the high levels of GDF15 observed in pregnant macaques could mediate paracrine effects in the placenta, we studied the expression levels of *GFRAL* mRNA in macaque placenta – the only known receptor for GDF15. However, *GFRAL* mRNA in the monkey placental tissue was below the detection limit (Ct > 35, data not shown).

**Figure 2.**
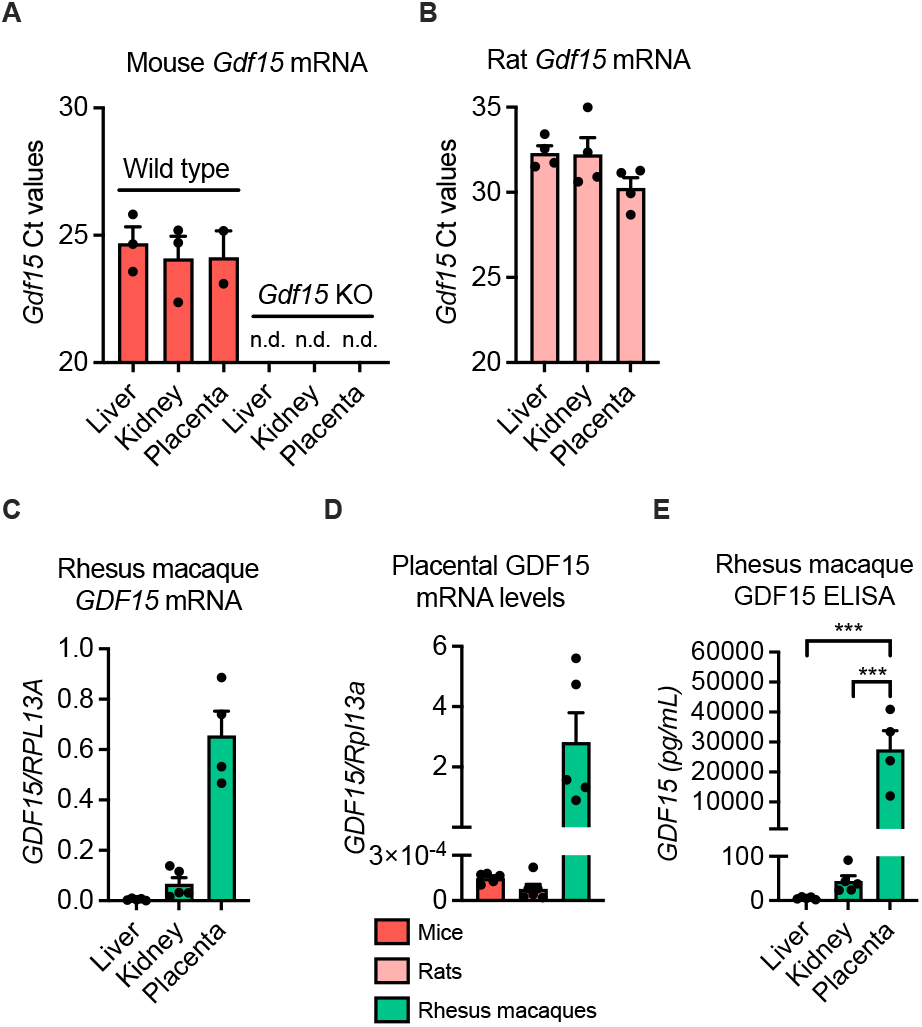
GDF15 tissue mRNA and protein during pregnancy. **A**. Mouse *Gdf15* mRNA levels in liver, kidney and placenta of WT (n=2-3) and GDF15 KO (n=3) mice. **B**. Rat *Gdf15* mRNA levels in liver, kidney and placenta (n=4). **C**. Monkey *GDF15* mRNA levels in liver (n=5), kidney (n=5) and placenta (n=4). **D**. Comparison of GDF15 mRNA levels across species, showing GDF15 expression in mice (n=5), rats (n=6), and macaques (n=5). **E**. GDF15 protein levels as measured by ELISA in liver (n=5), kidney (n=5), and placenta (n=4) from pregnant macaques. Data are presented as mean ± SEM. *** p < 0.001.

## Discussion

For decades, GDF15 has been recognized as one of the predominant hormones circulating during pregnancy^19^. Several studies have reported 50-200-fold increases in circulating GDF15 during pregnancy in humans^19,20,24^. This study corroborates these results and reveal that macaques display a similar increase in circulating GDF15 in response to pregnancy. Further, we demonstrate that in contrast to primates, rodents exhibit a very subtle increase in circulating GDF15 in response to pregnancy and thus might be unsuitable for studying the physiology of GDF15 in this condition. Together, these results indicate that non-human promates, but not rodents, are appropriate for studying the role of GDF15 in pregnancy and for evaluating GFRAL blocking agents as targets for prevention of morning sickness and HG.

Nausea and vomiting during pregnancy (NVP) is a very common disorder affecting up to 70% of pregnant women, although with a very high divergence in severity^26^. The underlying factors driving NVP are not known. Previously, human chorionic gonadotropin (hCG) has been main suspect given that its peak level in circulation coincide with the symptoms of NVP^25^. More recently, the GDF15-GFRAL pathway has been proposed as a likely mediator of nausea associated with pregnancy^23,24^. The placenta – at least in macaques - produces remarkably high levels of GDF15, contributing to plasma levels rising to 200-fold above baseline^18,19^. Considering the emetic role of exogenous GDF15 in preclinical models^14,16^, a causal connection between GDF15 and nausea and vomiting (morning sickness) during pregnancy is plausible. In support, Fejzo et al., reported that genetic variations of both *GDF15* and *GFRAL* are associated with NVP^23^. Furthermore, a correlation between GDF15 levels and the degree of morning sickness has been demonstrated^36^.

Several uncertainties regarding the physiological role of GDF15 during pregnancy remain. GDF15 levels in pregnant women continue to rise throughout the gestation period^20^, but in most pregnancies the morning sickness symptoms wear off after the first trimester^37^. Also, *all* pregnant women have very high circulating GDF15 levels, but ∼30% experience insignificant signs of nausea. From preclinical studies, it can be speculated that circulating GDF15 levels of >10,000 pg/ml are highly nauseating – but then how come all pregnant women are not sick with many emetic episodes? Whether this reflects interindividual variability to GDF15 exposure, e.g., via differences in GFRAL desensitization, has not been investigated. These uncertainties regarding the physiological role of GDF15 in pregnancy must be thoroughly clarified prior to administering novel GFRAL-blocking therapeutics to pregnant women. Could it be that GDF15 is essential for fetal development and/or that the primary role of elevated GDF15 during pregnancy is to ensure appropriate avoidance of teratogens? Or does GDF15 act in concert with other placental cytokines as immunomodulators to support a fetal-maternal tolerance?

In conclusion, we reveal that macaques exhibit a tremendous increase in placental and circulating GDF15 during pregnancy, comparable to what is reported in humans. In contrast, GDF15 is only negligible increased in pregnant mice and rats, questioning a physiological role for GDF15 in pregnancy in rodents. The species-specific differences in GDF15 biology suggest that strategies pursuing pharmacological interference of GDF15-GFRAL signaling should be translated with caution and preferably include preclinical testing in animal models with higher predictive value than rodents.

## Acknowledgements

We thank Charlotte Sashi Aier Svendsen for technical assistance. The Novo Nordisk Foundation Center for Basic Metabolic Research is an independent Research Center, based at the University of Copenhagen, Denmark, and partially funded by an unconditional donation from the Novo Nordisk Foundation (www.cbmr.ku.dk) (Grant number NNF18CC0034900).

